# Chronumental: time tree estimation from very large phylogenies

**DOI:** 10.1101/2021.10.27.465994

**Authors:** Theo Sanderson

## Abstract

Phylogenetic trees are an important tool for interpreting sequenced genomes, and their interrelationships. Estimating the date associated with each node of such a phylogeny creates a “time tree”, which can be especially useful for visualising and analysing evolution of organisms such as viruses. Several tools have been developed for time-tree estimation, but the sequencing explosion in response to the SARS-CoV-2 pandemic has created phylogenies so large as to prevent the application of these previous approaches to full datasets. Here we introduce Chronumental, a tool that can rapidly infer time trees from phylogenies featuring large numbers of nodes. Chronumental uses stochastic gradient descent to identify lengths of time for tree branches which maximise the evidence lower bound under a probabilistic model, implemented in a framework which can be compiled into XLA for rapid computation. We show that Chronumental scales to phylogenies featuring millions of nodes, with chronological predictions made in minutes, and is able to accurately predict the dates of nodes for which it is not provided with metadata.

## Introduction

The accumulation of mutations over time in living things means that genomes sequenced from a population capture information on the historical connections between its members. For viruses, these relationships can be usefully represented as a phylogenetic tree. The tips of such a tree are sequenced viruses, where we have a genome sequence, and typically also metadata on the date and location at which the sample was taken. Phylogenetic trees are often represented as a “distance-tree” in which the lengths of branches correspond to the genetic distance predicted between ancestral nodes and their descendants. However an alternative approach is to create a “time-tree”, where all nodes are positioned according to the date at which they are thought to have occurred. While establishing the dates of the tips is straightforward, using the metadata, inferring the likely dates for internal nodes requires the use of algorithms.

In recent years a number of approaches have been developed for creating time-trees. These include methods such as TreeTime (Sagulenko et al., 2018), TreeDater (Volz and Frost, 2017), BactDating (Didelot et al., 2018), wLogDate (Mai and Mirarab, 2020) and LSD (To et al., 2016), which all take as input a distance tree (and sometimes sequences) and use these to construct a predicted time tree. An alternative approach is to use BEAST (Suchard et al., 2018) to infer time and distance trees together from the sequences themselves. These algorithms can be assessed in two orthogonal dimensions. Firstly, how accurately do they model the underlying evolutionary dynamics? Longstanding MCMC-based approaches such as BEAST are likely to be the most preferable options if such fidelity is the only criterion on which an algorithm is being evaluated (Gill et al., 2020). However a second dimension has prompted the development of new tools: the ability of the algorithm to scale to datasets of interest, which are becoming increasingly large. BEAST-based analyses can take days or weeks for datasets with hundreds of sequences (Sagulenko et al., 2018), which prompted the development of the other approaches discussed. The unprecedented response to the SARS-CoV-2 pandemic has created a new magnitude of viral genomic data to which none of these tools easily scale. Nextstrain (Hadfield et al., 2018) has been an invaluable tool for analysis of viral genomic data during the SARS-CoV-2 pandemic, in part because it presents sequence data as easily interpretable time-trees, which are inferred using TreeTime. However these calculations are one of the key bottlenecks in the analysis, and together such bottlenecks mean that NextStrain analyses are typically limited to fewer than 10,000 sequences, sampled in a principled way. With more than 4,000,000 public SARS-CoV-2 genomes now sequenced, such analyses use only 0.25% of the available sequencing data (though sampled to provide as much information as possible). Importantly, however sophisticated a time-tree algorithm, its performance can still be limited by the data provided to it. Every node with chronological metadata creates a constraint on the time tree, providing information about the positions of unknown nodes. In the extreme case in which every circulating virus was sequenced every single day, inferring a time tree would be, in terms of logic, trivial. In countries that have been able to perform large scale genomic surveillance there have been times where a substantial portion (perhaps more than half) of all infections have been sequenced. Such dense data can provide so many constraints on the dates of internal nodes as to make inferring chronology simple, in terms of the sophistication of approach needed, even for a putative human curator working manually. What is not trivial is the scale of the data – constructing such a time tree for a dataset featuring millions of nodes and branches poses computational challenges, especially given the possibility of occasionally erroneous metadata. To our knowledge no existing tool is able to perform such an analysis.

Here we present Chronumental (“*chron*ologies from mon*umental* trees”), a tool that is able to quickly generate time trees from distance trees featuring millions of nodes. Chronumental represents the task of inferring a time tree as a series of matrix-based operations, allowing the use of efficient libraries recently developed for machine learning. It is capable of inferring a time tree from a tree of two million nodes in a matter of minutes on a consumer computer, and is able to tolerate the errors that inevitably occur in a subset of the metadata for very large datasets.

## Methods

### Algorithm

Let *r* be the root date *T*_*i*_ be the time-based branch length of the *i*th branch, *M*_*i*_ be the distance-based length of the *i*th branch, *μ* be the mutation rate, *Di* be the observed date of the *i*th terminal node, *Ei* be the error associated with the *i*th terminal node (determined by whether date takes the form yyyy-mm-dd, yyyy-mm or yyyy), and 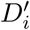 be the calculated date of the *i*th terminal node.

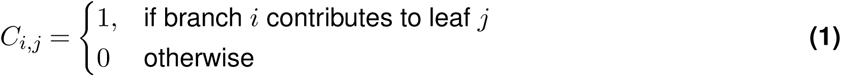

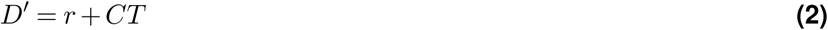

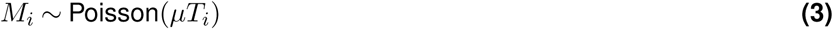

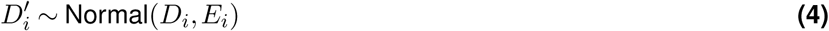

### The Chronumental algorithm

Chronumental is implemented in Python using NumPyro (Phan et al., 2019), a probabilistic programming library built on top of JAX (Bradbury et al., 2018), a system for compiling differentiable matrix operations for efficient execution using XLA (a compiler for linear algebra”). In essence we build a semi-probabilistic model of the underlying dynamics, with unknown quantities as latent variables, then perform maximum *a priori* estimation to maximise the evidence lower bound by stochastic gradient descent. That is to say, we take a series of small steps in the values of the latent variables, which are expected (by differentiation) to gradually increase the correspondence between the model and the observed data.

The input to our algorithm is a rooted tree, with branch lengths measured in terms of numbers of mutations. For most of the tips of the tree, there will be associated date metadata. Our aim is to estimate the length of time represented by each edge of the tree, and hence the dates associated with all nodes, including the internal nodes that lack metadata. We treat the length of each branch in units of time as a learnable parameter (which must be >= 0).

From these time-lengths we can calculate a date for each tip node, by representing its date as the sum of the time-lengths of the edges leading up to it, added to the date of the root (which is treated as a further learnt parameter). An important insight was that we could represent the summation of branch lengths to estimate node dates as a notional matrix multiplication, by imagining constructing a vast matrix in which one dimension represents the leaf nodes, and one dimension represents the internal branches, with a 1 at each element *x*_*i,j*_ where branch j contributes towards the date of leaf node i, and a 0 where it does not. When this matrix is multiplied by a vector of time-lengths for each branch it would yield the date corresponding to each leaf node^1^. Such a matrix would contain *>* 10^12^ elements, dwarfing any resources, but since almost all elements are 0s, it can be represented as a “sparse matrix”, encoded in coordinate list (COO) format, with the matrix multiplication performed through ‘take’ and ‘segment_sum’ XLA operations. Representing the operations in this way allows them to be efficiently compiled in XLA, which creates a differentiable graph of arithmetic operations.

We treat these modelled final dates as the centres of normal distributions, with observations corresponding to the dates actually seen. Notionally the variance in this normal distribution has two sources: firstly general additional sampling dynamics which aren’t modelled and so appear as noise, and secondly gross metadata errors. Treating these observations as samples of a random distribution permits Chronumental to occasionally place samples very differently in time from where their metadata would suggest, which is essential given that some samples will have metadata errors that would otherwise provide such a strong constraint as to prevent a reasonable time tree being created. Additionally, Chronumental is able to accept dates at a range of precisions, from days (2021-03-05) to months (2021-03) to years (2021). The variance of the normal distributions for the tip dates is scaled according to the indicated uncertainty.

The second set of data available to our algorithm is the number of mutations that occurred in each branch. We consider these to be observations of Poisson distributions whose rates are calculated by multiplying the time-length of each branch by a learnt parameter representing the mutation rate (treated as the same for all branches). This aspect of the model means that, within the constraints above, branch lengths in distance are made to correlate with branch lengths in time. The starting value of the clock rate can be set by the user, or the clock rate can be entirely fixed at a manually given value. If the user does neither, the initial value of the clock rate is automatically estimated by root-to-tip regression.

The model is fit by using the ‘Stochastic Variational Inference’ module of NumPyro. The Adam optimiser is used to adjust the latent variables to maximise the evidence lower bound. Although this approach uses the variational inference module, we do not aim to estimate the uncertainty in our predictions of branch time lengths, or in node dates. In the guide for the model, branches’ time-lengths are represented as Delta distributions with a single value. We do provide the optional ability to model uncertainty in the mutation rate, though in any given sample of the model this is treated as the same across all branches.

Chronumental uses TreeSwift (Moshiri, 2020) to read and manipulate trees rapidly. The command-line parameters are inspired in large part by TreeTime. Chronumental is open-source, with code available at github.com/theosanderson/chronumental.

### Dataset and tree fitting

Our immediate motivation in developing Chronumental was to allow time trees to be constructed for the very large phylogenetic trees generated during the SARS-CoV-2 pandemic. An open-source repository, sarscov2phylo (Lanfield, 2020), was an initiative that created large public phylogenies from GISAID data until November 2020. The development of UShER (Turakhia et al., 2021), to permit rapid expansion of such a tree by sequential addition of new samples by maximum parsimony, enabled phylogenetics to keep up with the ever-expanding sequencing efforts that have occurred during 2021. There are two major such trees: a public tree maintained maintained by researchers at UCSC (McBroome et al., 2021), which uses data available without legal restrictions from the INSDC databases (Arita et al., 2021), COG-UK (Nicholls et al., 2021), and the database of the China Center for Bioinformation; and the Audacity tree maintained within the GISAID Initiative (Elbe and Buckland-Merrett, 2017). Both groups maintain convenient metadata sets for the associated datasets.

We used Chronumental to create time trees for both of these trees, using default parameters other than increasing the number of steps to 2000. In the interests of reproducible open-source analysis we focus our benchmarking studies on the UCSC public tree. Its creators maintain an archive of trees from various points in time, and here we used the 2021-09-15 tree.

### Prediction of dates for terminal nodes without metadata

We initially assessed the general plausibility of our trees by visualising and exploring it in Taxonium (Sanderson, 2022). For a more quantitative assessment, we conducted experiments in which we blinded the algorithm to some of the available date metadata. While the nodes whose dates must be inferred in time-tree estimation are typically the tree’s internal nodes, the algorithm is equally able to estimate the date of any uncertain tip nodes (which may arise even in real applications, where some metadata is missing). Estimating the date of a tip node essentially requires estimation of the date of an internal node, and also then estimation of the length of time between that internal node and the tip. Therefore, by blinding the algorithm to dates for a certain number of tips we can assess how well it recapitulates the ground-truth, providing an upper bound on the error with which it estimates the dates of internal nodes.

We performed such an analysis on the 2021-09-15 public tree, with a wide range of proportions of the metadata blinded to assess how important densely sampled data are to predicting node dates with this approach.

### Speed and memory usage comparisons

To provide a sense of the challenge that Chronumental was designed to address, we compared its running times and memory usage with those from an existing tool, TreeTime, for a range of tree sizes. We started from the 2021-09-31 public tree, and used gotree’s `prune` function (Lemoine and Gascuel, 2021) to retain a small proportion of nodes, which we increased in increments. We then predicted time trees with both Chronumental, TreeTime (running with simply the `--dates`, `--tree`, `--keep-root` and `--sequence_length` parameters)stopping once runtime reached 100 minutes. Chronumental was run for 1000 steps in all cases, either in CPU mode or in GPU mode. We included TreeTime as perhaps the most-used specialist tree dating tool (at least based on citations). Previous benchmarking results suggest longer runtimes from tools such as BEAST and wLogDate (Mai and Mirarab, 2020).

### Comparing outputs to a traditional algorithm

To compare our algorithm to one previously used, we used the dataset of Ebola genomes from the 2014 outbreak of Ebola in West Africa (Dudas et al., 2017), using the 350 genomes and metadata presented in the treetime_examples repository by Sagulenko et al. (2018). We firstly ran TreeTime, with the `--confidence` and `--covariance` parameters (and providing the sequence alignment). In the course of this analysis TreeTime re-rooted the distance tree, which it output alongside the time tree. We used this re-rooted distance tree as an input to Chronumental, along with the metadata, and compared its results for the internal nodes lacking metadata to those obtained with TreeTime.

### Method comparison: simulated data

We also tested the ability of Chronumental to predict the internal dates of nodes, using simulated datasets from To et al. (2016) which has been reanalysed by a number of papers since. This paper provided four types of simulated tree, each with 100 replicates ^2^. We downloaded the PhyML distance trees from the strict molecular clock model. We downloaded the corresponding true time trees from a repository from Mai and Mirarab (2020) ^3^. We calculated time trees using Chronumental, LSD2, and TreeTime and calculated the errors between each algorithm’s assessment of branch length in time and the true value.

In a second analysis, we randomly shuffled date metadata for 5% of tips to study different algorithms’ ability to tolerate this erroneous metadata.

## Results

### Time trees with millions of nodes

We used Chronumental to assign dates to each node in the 2019-09-15 tree created by UCSC (Figure 1). Within the first 180 steps of fitting, the algorithm was able to place the median terminal node within a day of its position in the metadata. After the algorithm completed, 90% of nodes were placed within 3 days of their metadata position (and 99% within 2 weeks). Supervision is provided on these dates, so this simply measures the algorithm’s ability to reconcile date metadata into a tree structure, rather than its ability to predict the dates for nodes where the date is unknown.

**Figure 1.**
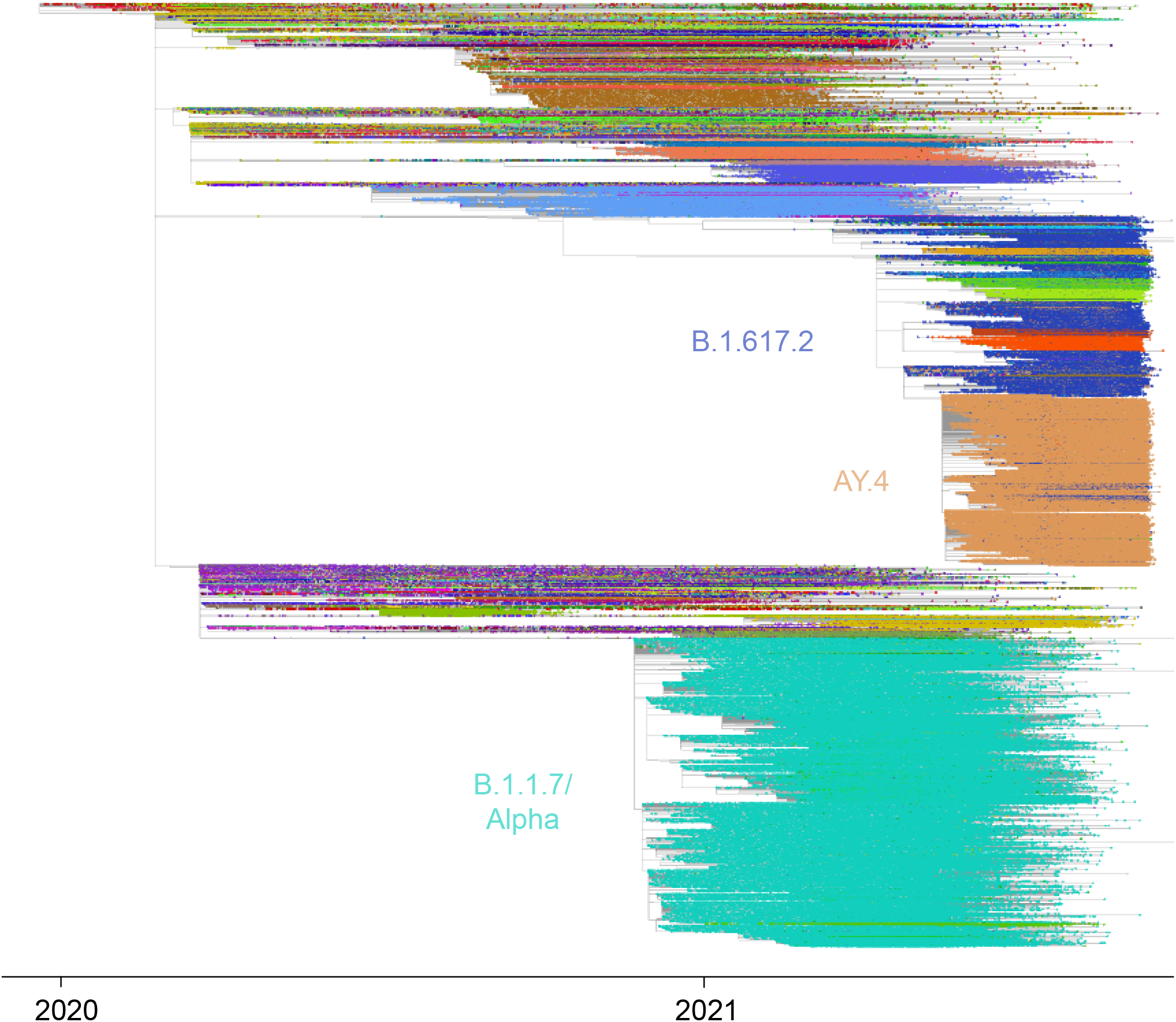
A time tree calculated using Chronumental for a phylogeny of 2,045,884 nodes. The starting point for this chronology is the 2021-09-15 UCSC public distance tree (McBroome et al., 2021). This visualisation is from Taxonium, with points filtered to reduce overplotting.

Functionality for inferring time trees with Chronumental is now built into Taxonium (Sanderson, 2022), and the latest time trees for public data can be visualised at http://Cov2Tree.org, where the Chronumental output is available for download.

### Identification of anachronistic nodes

Though the vast majority of points were assigned dates very close to their dates recorded in the metadata, we found 182 nodes were placed more than 90 days away from where their metadata indicated. In doing this the algorithm incurs a large cost to its loss function (a true date so far from the observed date is considered highly unlikely), and so the expectation is that this will only occur where placing that node close to the date recorded in its metadata is also extremely unlikely, given the mutation profile and tree topology observed. To consider these possibilities, we plotted the metadata date of sequences against their observed date (Figure 2). We see, as expected, an extremely close relationship, but with rare outliers. By categorising nodes according to the lineage of their sequences, we can see whether the genotypes of the samples plausibly correspond to their metadata date. We found that outlier samples in general belonged to lineages that are known not to have existed at the time at which their metadata would indicate, suggesting that the metadata is inaccurate for these sequences, and the calculated date significantly more correct. This means that Chronumental analyses are able to identify data quality issues in a similar way to tools such as TempEst (Rambaut et al., 2016) and TreeTime.

**Figure 2.**
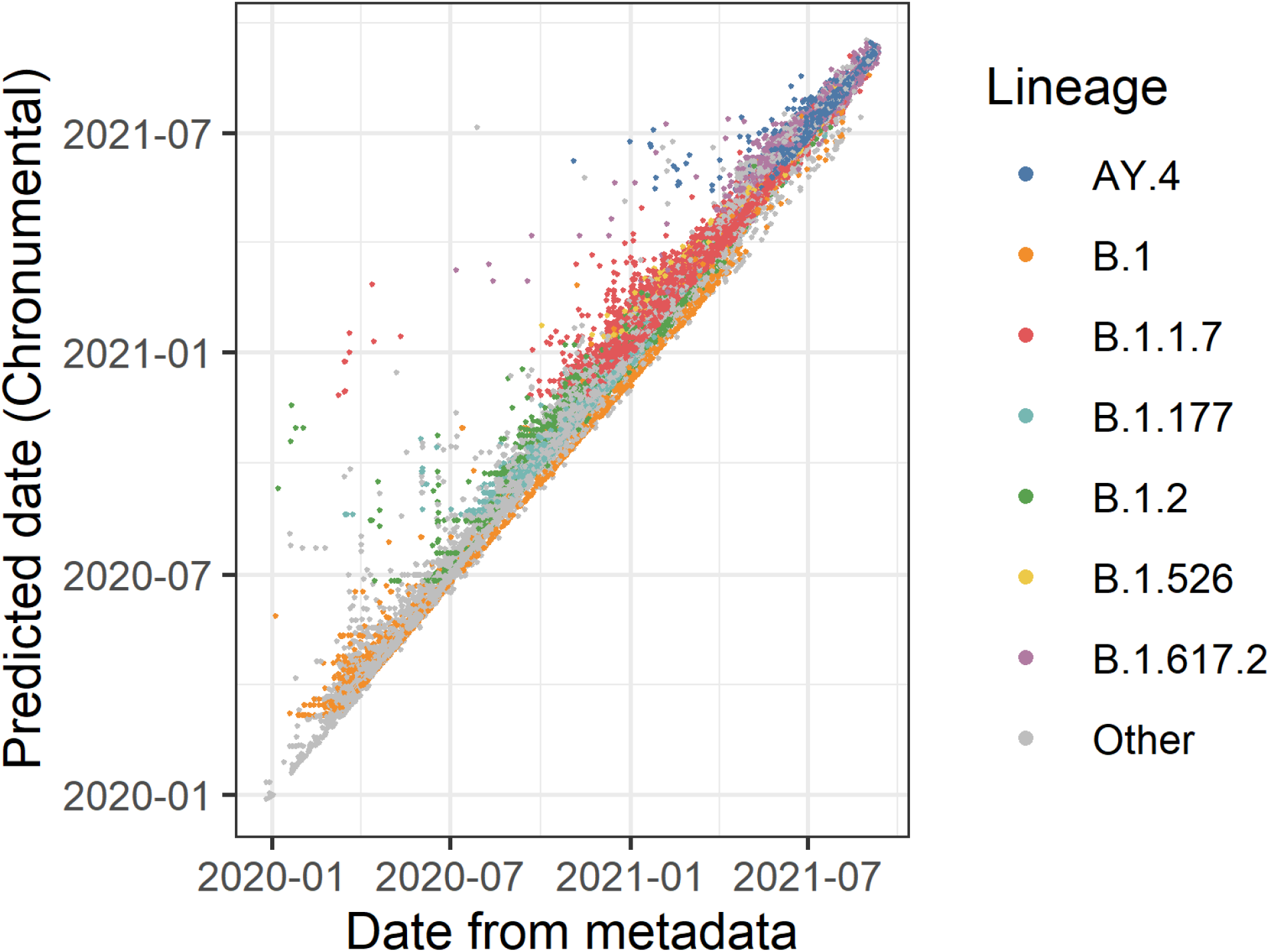
Nodes placed unexpectedly in time by Chronumental appear to have incorrect metadata. This graph plots supposed ground-truth dates from metadata against Chronumental’s predicted dates. While there is general agreement, in rare cases of discordance it appears from the genotype that Chronumental may be more correct, as in many cases discordant points are lineages suggested to have occurred before their emergence.

We happened to perform the same analysis on a later dataset (2021-10-09) in which due to a temporary metadata error, a relatively large set of sequences had been given erroneous dates. Again, such sequences were immediately apparent on a plot. Due perhaps to the large number of such sequences, predicted dates for some sequences lay somewhere between their date as indicated in the metadata and the actual likely ground truth date (late due to the presence of the Delta lineage). This suggests that the most robust approach would be an iterative one in which an initial time tree is fit, and used to identify spurious metadata, which is then excluded in a subsequent analysis. It is also possible that one could improve the approach by using a spike and slab prior for the distribution of observed dates – a mixture of a very tight normal distribution representing samples with correct metadata and a high-variance, or even uniform, distribution representing occasional metadata errors – but this has implementation complexities in NumPyro.

To facilitate analyses of anachronistic sequences, or the imputing of dates for sequences with missing metadata (which can be either wholly missing, or coarse to the level of months or years – creating a rough prior), Chronumental provides an option to export a TSV file containing its own calculated dates for all tips on the tree.

### Assessing predictive performance using suppressed metadata

To establish the predictive performance of Chronumental, we created a series of datasets in which we hid from the algorithm metadata for a subset of tips, from 10% up to 99.95%. We then assessed the error in prediction for these nodes as compared to the known ground truth (Figure 3). At minimum predicting such a tip requires estimating the date of an internal node, and then estimating the branch length to add to it. In more extreme cases many tips would have required estimating multiple branch lengths.

**Figure 3.**
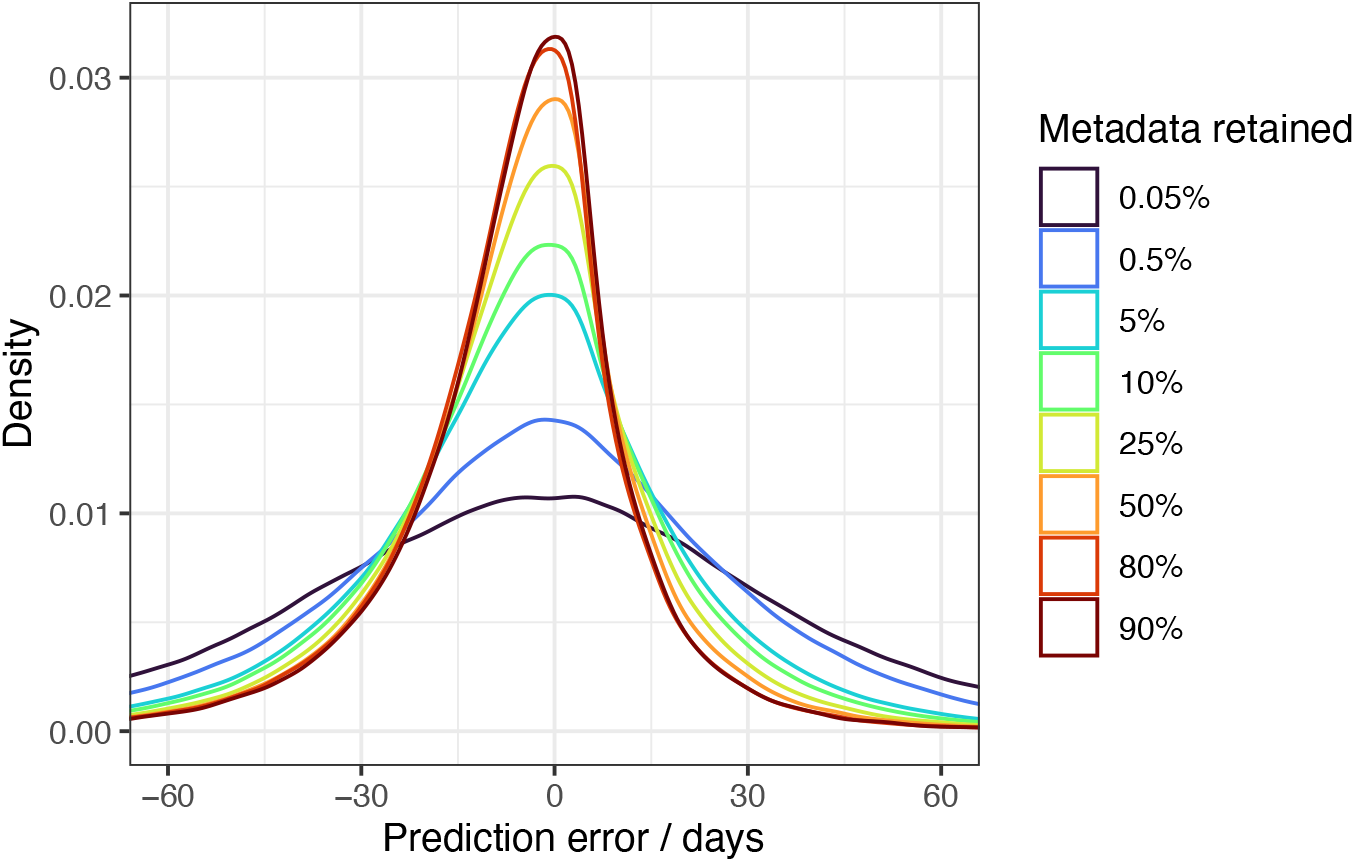
Prediction of ground truth for nodes with metadata hidden for Chronumental in a 2 million node tree of SARS-CoV-2 genomes. Colours represent the proportion of nodes that were suppressed. In all cases only data from suppressed nodes is plotted.

In the 10%-suppressed setting, which provides an indication of the bounds on our ability to predict dates for internal nodes, 89% of predictions were within 30 days of their known ground truth value. In the 90%-suppressed setting, providing a sense of a more sparsely sampled tree, still 81% of predictions were within 30 days of ground-truth. When the amount of dates retained fell to just 0.005% (99.95%-suppressed), 54% of dates could be predicted within 30 days.

### Comparing predictions to those for a traditional algorithm for the well-studied West African Ebola virus epidemic dataset

To provide a direct comparison to an existing dataset, we used Chronumental to predict dates for 350 sequences from the 2014-16 West Africa Ebolan Ebola virus epidemic (Dudas et al., 2017), using a dataset prepared and previously analysed by Sagulenko et al. (2018). We applied both TreeTime (which can generate confidence intervals) and Chronumental to this dataset. We note that this is not a situation in which we would recommend the use of Chronumental – tools designed to optimally analyse these smaller datasets will likely yield better results. We plotted the dates predicted for internal nodes by the two methods against each other (Figure 4). We found that 72% of the dates predicted by Chronumental lay within the confidence intervals calculated by TreeTime.

**Figure 4.**
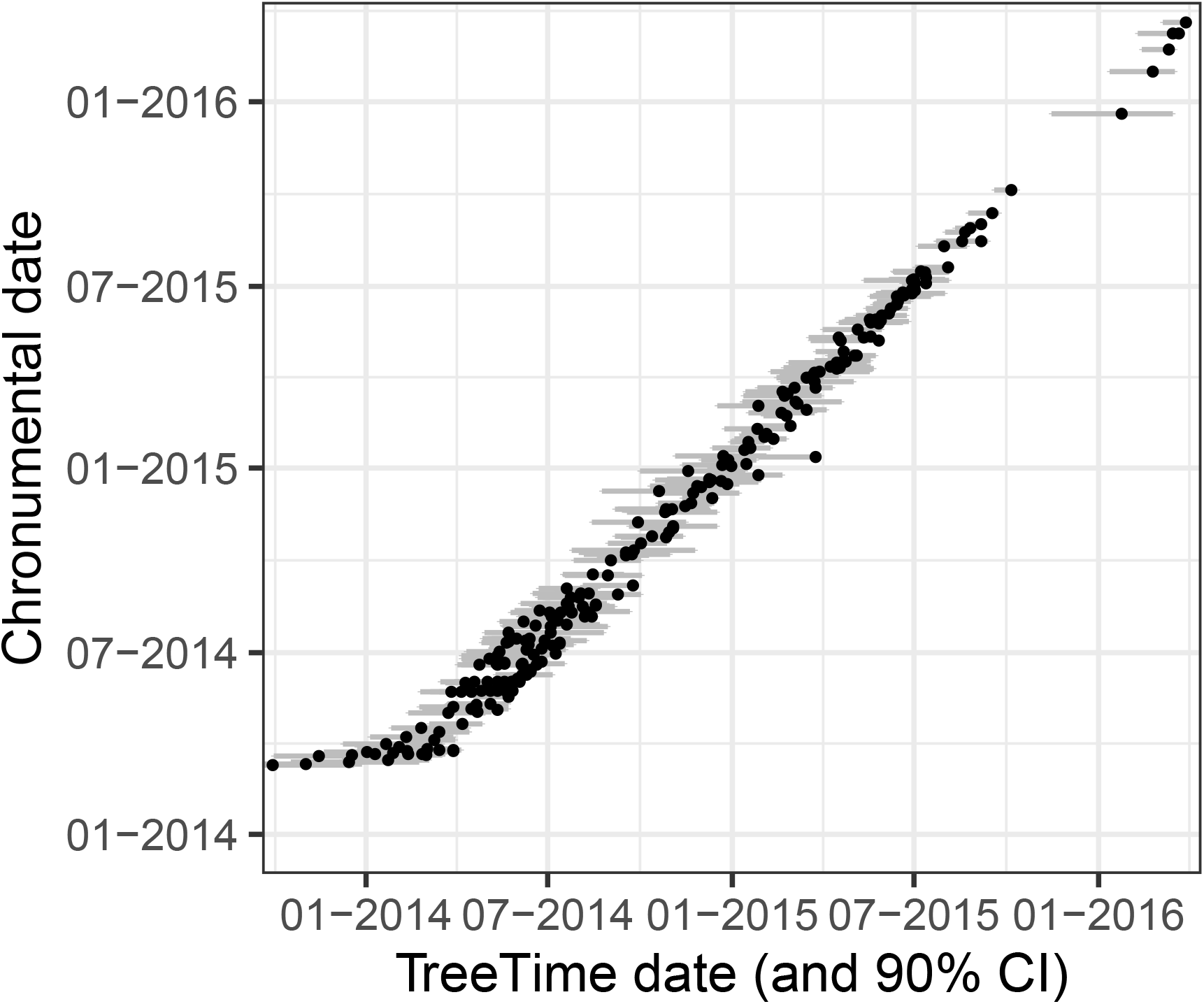
Comparison of dates predicted by TreeTime (x) and Chronumental (y) for the internal nodes of a 350 sequence phylogeny from the 2014 West African Ebola virus epidemic. Lines indicate 90% confidence intervals from TreeTime.

### Speed and memory usage comparison

To illustrate the problem Chronumental attempts to solve, we compared the runtime and memory usage of Chronumentaland TreeTime for a range of differently sized phylogenies (Figure 5). Compared to TreeTime, running time could be perhaps a hundredfold lower with Chronumental at large tree sizes, with resource usage at least an order of magnitude lower. However such comparisons are fraught with caveats. Chronumental’s runtime varies with the number of steps chosen, depending on the level of precision required, and a more experienced user of TreeTime might also customise it for increased performance. These results should be interpreted as general trends, and in the context that TreeTime provides the potential for carrying out many other features. We also compared Chronumental running in GPU mode and CPU mode. We found that as tree size approached a million nodes, running on a GPU prevented runtime from increasing substantially, but that prior to this running on a CPU was preferable.

**Figure 5.**
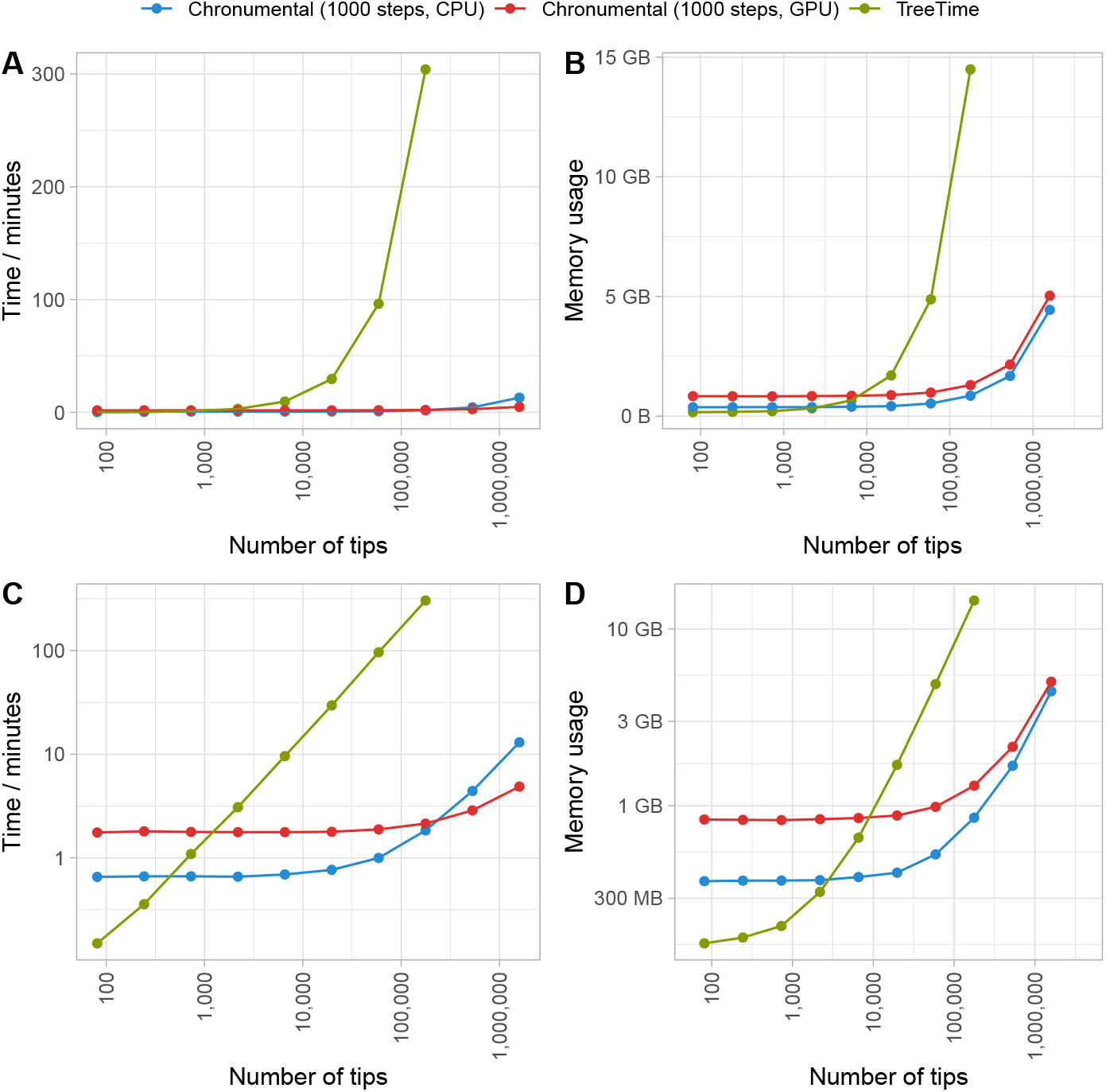
Runtime (A,C) and memory requirements (B,D) for TreeTime and Chronumental for a range of tree sizes, plotted on linear (A,B) and logarithmic (C,D) scales.

## Discussion

We have shown that Chronumental can rapidly infer time trees from phylogenies featuring millions of nodes. Examining the algorithm’s output when a subset of input metadata are suppressed suggests strong predictive performance, and application to a classic dataset largely recapitulates the results of an alternative approach. Chronumental’s ability to provide chronologies for large phylogenies unlocks new possibilities for visualisation and analyses of complete genomic datasets.

Our tool has been developed to tackle a specific problem – very large trees – unserved by existing approaches. Chronumental does not feature all of the features offered by some alternative tools. There is currently no capacity for polytomy resolution. An algorithm could be developed that would take a time tree previously optimised by Chronumental and identify the most likely resolutions of each polytomy, given the date estimates from Chronumental, and perhaps geographical metadata – running alternately with Chronumental date updates. Chronumental is not currently able to re-root a tree, and must be supplied with a rooted tree. In the case of SARS-CoV-2, rooting with an old member of the A lineage is relatively straightforward.

Chronumental does not identify ancestral residues at genomic positions as does TreeTime. UShER (Turakhia et al., 2021) can efficiently perform ancestral state reconstruction for very large trees, and allows sequential addition of samples to make these trees larger still. The combination of UShER (with initial use of iqtree (Minh et al., 2020) or other tools to create an initial tree) and Chronumental, goes some way towards bringing some of the powerful analyses enabled by NextStrain Augur (Hadfield et al., 2018) to entire sequence datasets with millions of sequences, and we are in parallel developing tools (Sanderson, 2022) that allow such datasets to be visualised.

Chronumental’s statistical approach may offer fewer guarantees than some other tools, and does not measure uncertainty, nor nucleotide-specific substitution rates. In particular, this could pose issues for parts of the very large tree where samples are very sparse, such as countries with low current capacities for genomic surveillance. Chronumental currently offers only a strict clock model, rather than allowing rates of evolution to differ between branches of a tree. We can see a path for adapting the approach to use a correlated relaxed clock model. Chronumental is likely to be best suited to analyses of densely sampled trees across short periods of time. We would recommend the use of tools such as TreeTime or TreeDater wherever datasets are small enough to permit this.

Millions of people acquire infectious diseases each day, and the proportion of these cases that are genome-sequenced is likely to rise over time. This new scale of data collection will advance our understanding of transmission dynamics, but also pose new challenges for analytic workflows. Chronumental, and future work building on it, provides a contribution to a new scalable infrastructure for genomic epidemiology.

## Reproducibility

Chronumental is open-source, with code available at github.com/theosanderson/chronumental. Our code for performing the analyses in this paper is available at github.com/theosanderson/chron_analysis.

## Acknowledgements

We thank Alex Kramer and Russell Corbett-Detig for useful discussions, and Angie Hinrichs for maintaining the UCSC tree. We are grateful to all who have submitted sequences to the INSDC and GISAID databases, and to the databases’ curators. We thank Ricardo Henriques and Steve Royle for the LaTeX template used for this manuscript.

TS is funded by a fellowship [210918/Z/18/Z] from the Wellcome Trust. This work was supported by the Francis Crick Institute, which receives its core funding from Cancer Research UK [FC01121], the UK Medical Research Council [FC01121], and the Wellcome Trust [FC01121]. For the purpose of Open Access, the authors have applied a CC BY public copyright licence to any Author Accepted Manuscript (AAM) version arising from this work.

## Footnotes

1 In review it was pointed out that To et al. (2016) made use of a similar matrix representation much earlier

2 http://www.atgc-montpellier.fr/LSD/

3 https://github.com/uym2/LogDate-paper/tree/master/simulated_data/True_time_trees

## Bibliography

Arita, M., Karsch-Mizrachi, I., and Cochrane, G. The international nucleotide sequence database collaboration. Nucleic Acids Res., 49(D1):D121–D124, Jan. 2021.

Bradbury, J., Frostig, R., Hawkins, P., Johnson, M. J., Leary, C., Maclaurin, D., Necula, G., Paszke, A., VanderPlas, J., Wanderman-Milne, S., and Zhang, Q. JAX: composable transformations of Python+NumPy programs, 2018. URL http://github.com/google/jax.

Didelot, X., Croucher, N. J., Bentley, S. D., Harris, S. R., and Wilson, D. J. Bayesian inference of ancestral dates on bacterial phylogenetic trees. Nucleic Acids Res., 46(22):e134, Dec. 2018.

Dudas, G., Carvalho, L. M., Bedford, T., Tatem, A. J., Baele, G., Faria, N. R., Park, D. J., Ladner, J. T., Arias, A., Asogun, D., Bielejec, F., Caddy, S. L., Cotten, M., D’Ambrozio, J., Dellicour, S., Di Caro, A., Diclaro, J. W., Duraffour, S., Elmore, M. J., Fakoli, L. S., Faye, O., Gilbert, M. L., Gevao, S. M., Gire, S., Gladden-Young, A., Gnirke, A., Goba, A., Grant, D. S., Haagmans, B. L., Hiscox, J. A., Jah, U., Kugelman, J. R., Liu, D., Lu, J., Malboeuf, C. M., Mate, S., Matthews, D. A., Matranga, C. B., Meredith, L. W., Qu, J., Quick, J., Pas, S. D., Phan, M. V. T., Pollakis, G., Reusken, C. B., Sanchez-Lockhart, M., Schaffner, S. F., Schieffelin, J. S., Sealfon, R. S., Simon-Loriere, E., Smits, S. L., Stoecker, K., Thorne, L., Tobin, E. A., Vandi, M. A., Watson, S. J., West, K., Whitmer, S., Wiley, M. R., Winnicki, S. M., Wohl, S., Wölfel, R., Yozwiak, N. L., Andersen, K. G., Blyden, S. O., Bolay, F., Carroll, M. W., Dahn, B., Diallo, B., Formenty, P., Fraser, C., Gao, G. F., Garry, R. F., Goodfellow, I., Günther, S., Happi, C. T., Holmes, E. C., Kargbo, B., Keïta, S., Kellam, P., Koopmans, M. P. G., Kuhn, J. H., Loman, N. J., Magassouba, N., Naidoo, D., Nichol, S. T., Nyenswah, T., Palacios, G., Pybus, O. G., Sabeti, P. C., Sall, A., Ströher, U., Wurie, I., Suchard, M. A., Lemey, P., and Rambaut, A. Virus genomes reveal factors that spread and sustained the ebola epidemic. Nature, 544(7650):309–315, Apr. 2017.

Elbe, S. and Buckland-Merrett, G. Data, disease and diplomacy: GISAID’s innovative contribution to global health. 1(1):33–46, Jan. 2017. doi: 10.1002/gch2.1018.

Gill, M. S., Lemey, P., Suchard, M. A., Rambaut, A., and Baele, G. Online bayesian phylodynamic inference in BEAST with application to epidemic reconstruction. Molecular Biology and Evolution, 37:1832–1842, June 2020. doi: 10.1093/molbev/msaa047. Published: 26 February 2020.

Hadfield, J., Megill, C., Bell, S. M., Huddleston, J., Potter, B., Callender, C., Sagulenko, P., Bedford, T., and Neher, R. A. Nextstrain: real-time tracking of pathogen evolution, 2018.

Lanfield, R. roblanf/sarscov2phylo: 13-11-20, 2020. URL https://zenodo.org/record/3958883.

Lemoine, F. and Gascuel, O. Gotree/goalign: toolkit and go API to facilitate the development of phylogenetic workflows. 3(3), June 2021. doi: 10.1093/nargab/lqab075.

Mai, U. and Mirarab, S. Log Transformation Improves Dating of Phylogenies. Molecular Biology and Evolution, 38(3):1151–1167, sep 4 2020.

McBroome, J., Thornlow, B., Hinrichs, A. S., Kramer, A., De Maio, N., Goldman, N., Haussler, D., Corbett-Detig, R., and Turakhia, Y. A daily-updated database and tools for comprehensive SARS-CoV-2 mutation-annotated trees. Mol. Biol. Evol., Sept. 2021.

Minh, B. Q., Schmidt, H. A., Chernomor, O., Schrempf, D., Woodhams, M. D., von Haeseler, A., and Lanfear, R. IQ-TREE 2: New models and efficient methods for phylogenetic inference in the genomic era. Mol. Biol. Evol., 37(5):1530–1534, May 2020.

Moshiri, N. Treeswift: A massively scalable python tree package. SoftwareX, 11:100436, 2020. doi: 10.1016/j.softx.2020.100436.

Nicholls, S. M., Poplawski, R., Bull, M. J., Underwood, A., Chapman, M., Abu-Dahab, K., Taylor, B., Colquhoun, R. M., Rowe, W. P. M., Jackson, B., Hill, V., O’Toole, Á., Rey, S., Southgate, J., Amato, R., Livett, R., Gonçalves, S., Harrison, E. M., Peacock, S. J., Aanensen, D. M., Rambaut, A., Connor, T. R., Loman, N. J., and COVID-19 Genomics UK (COG-UK) Consortium. CLIMB-COVID: continuous integration supporting decentralised sequencing for SARS-CoV-2 genomic surveillance. Genome Biol., 22(1):196, July 2021.

Phan, D., Pradhan, N., and Jankowiak, M. Composable effects for flexible and accelerated probabilistic programming in numpyro, 2019.

Rambaut, A., Lam, T. T., Max Carvalho, L., and Pybus, O. G. Exploring the temporal structure of heterochronous sequences using tempest (formerly path-o-gen). Virus evolution, 2(1):vew007, 2016.

Sagulenko, P., Puller, V., and Neher, R. A. TreeTime: Maximum-likelihood phylodynamic analysis. Virus Evol, 4(1):vex042, Jan. 2018.

Sanderson, T. Taxonium: a web-based tool for exploring large phylogenetic trees. bioRxiv, 2022.

Suchard, M. A., Lemey, P., Baele, G., Ayres, D. L., Drummond, A. J., and Rambaut, A. Bayesian phylogenetic and phylodynamic data integration using BEAST 1.10. Virus Evol, 4(1):vey016, Jan. 2018.

To, T.-H., Jung, M., Lycett, S., and Gascuel, O. Fast dating using Least-Squares criteria and algorithms. Syst. Biol., 65(1):82–97, Jan. 2016.

Turakhia, Y., Thornlow, B., Hinrichs, A. S., De Maio, N., Gozashti, L., Lanfear, R., Haussler, D., and Corbett-Detig, R. Ultrafast sample placement on existing trees (UShER) enables real-time phylogenetics for the SARS-CoV-2 pandemic. Nat. Genet., 53 (6):809–816, June 2021.

Volz, E. M. and Frost, S. D. W. Scalable relaxed clock phylogenetic dating, 2017.

